# SimReadUntil for Benchmarking Selective Sequencing Algorithms on ONT Devices

**DOI:** 10.1101/2023.11.01.565133

**Authors:** Maximilian Mordig, Gunnar Rätsch, André Kahles

## Abstract

**Motivation:** The Oxford Nanopore Technologies (ONT) ReadUntil API enables selective sequencing, which aims to reduce time spent on sequencing uninteresting reads in favor of more interesting reads, e.g., to deplete or enrich certain genomic regions. The performance gain depends on the selective sequencing decision-making algorithm (SSDA) which decides whether to reject a read, stop receiving a read or wait for more data. Since real runs are time-consuming and costly (at scale), simulating the ONT device with support for the ReadUntil API is highly beneficial to compare and optimize the parameters of SSDAs. Existing software like MinKNOW and UNCALLED only return raw signal data, are memory-intensive, require huge and often unavailable multi-fast5 files (*≥* 100GB) and are not clearly documented.

**Results:** We present the ONT device simulator *SimReadUntil* that takes a set of full (real or simulated) reads as input, distributes them to channels and plays them back in real time including mux scans, channel gaps and blockages, and allows to unblock (reject) reads as well as stop receiving data from them (imitating the ReadUntil API). Our modified ReadUntil API provides the basecalled reads rather than the raw signal to reduce computational load and focus on the SSDA rather than basecalling. Tuning the parameters of tools like ReadFish and ReadBouncer becomes easier because no GPU is required anymore for basecalling. We offer various methods to extract simulation parameters from a sequencing summary file and compare them. *SimReadUntil* ‘s gRPC interface allows standardized interaction with a wide range of programming languages.

**Availability:** The code is freely available on GitHub (https://github.com/ratschlab/sim_read_until) along with a fully worked use case that combines the simulator with ReadFish (and optionally NanoSim).

**Supplementary information:** Supplementary data are available at Bioinformatics online.

## 1. Introduction

Nanopore sequencing has emerged as a promising third-generation sequencing technology that delivers long reads at an affordable price. An Oxford Nanopore Technologies (ONT) device consists of a flowcell, which has a set of channels that sequence reads in parallel. In a single channel, a read is followed by a read gap, which is the time for a new read to move into the pore. Reads interact biologically with the pore which deteriorates the pore over time and results in longer read gaps. After 72h of sequencing, the flowcell usually needs to be replaced (costing around 500$). Unlike other sequencing technologies, ONT devices support the ReadUntil API (Loose et al., 2016) which gives real-time access to the read in-progress and allows to perform actions on the sequencer: reject a read, stop receiving data from a read but keep sequencing it (to reduce processing overhead), or no action to wait for more data from a read (by default). *Selective sequencing* / *adaptive sampling* leverages the ReadUntil API to enrich target sequences or deplete non-target sequences by delegating the decisions on which actions to perform to a selective sequencing decision-making algorithm (SSDA). An SSDA typically has various tunable parameters, e.g., the trade-off between a high-accuracy (compute-intensive) and a low-accuracy (fast) basecaller, the number of basepairs to take into account for read mapping, the number of mapping attempts, or the hyperparameters of a learning algorithm. Simulating the ONT device can help with tuning the SSDA parameters by reducing sequencing cost and accelerating the parameter optimization through parallel execution.

Existing simulators focus on generating reads at the raw signal level (Li et al., 2018) or nucleotide level (Yang et al., 2017). They do not respond to selective sequencing decisions and do not model the read gaps or reduced pore throughput due to pore blockage. The only available ONT simulators with ReadUntil support are ONT’s for-research MinKNOW controller software (Oxford Nanopore Technologies Ltd.) and UNCALLED (Kovaka et al., 2021). They do not permit to effectively evaluate SSDAs because they are memory-hungry and require raw signals which are both huge and often unavailable. Table 1 and supplementary Figure 1 give a detailed comparison of existing simulators. The concurrent work Icarust (Munro et al., 2023) is an implementation in Rust related to ours, but returns raw signal level data rather than basecalled data. It does not extract parameters from an existing run, assumes constant gaps between reads, does not offer acceleration and mux scans, and simulates perfect reads only.

**Table 1.**
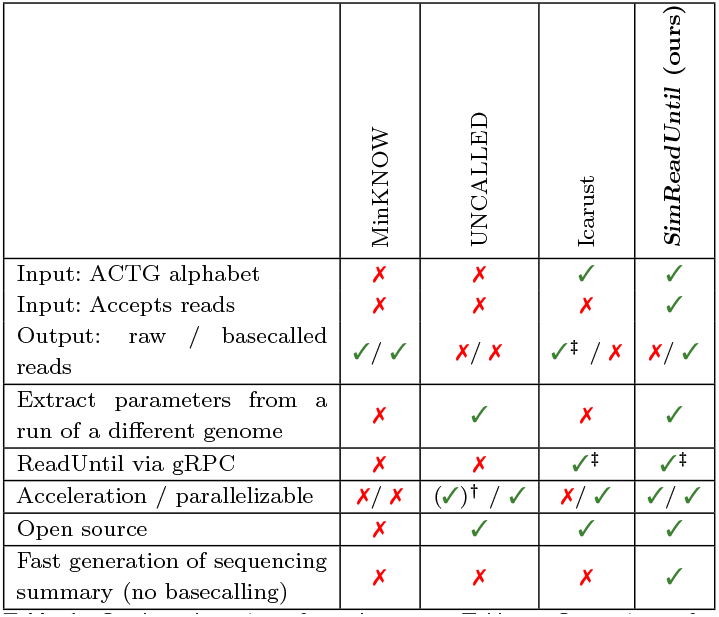
Condensed version of supplementary Table 2. Comparison of simulators with ReadUntil functionality. Icarust was not yet officially published, so it may still change.

|  | MinKNOW | UNCALLED | Icarust | SimReadUntil (ours) |
| --- | --- | --- | --- | --- |
| Input: ACTG alphabet | ✗ | ✗ | ✓ | ✓ |
| Input: Accepts reads | ✗ | ✗ | ✗ | ✓ |
| Output: raw / basecalled reads | ✓ / ✓ | ✗ / ✗ | ✓<sup>‡</sup> / ✗ | ✗ / ✓ |
| Extract parameters from a run of a different genome | ✗ | ✓ | ✗ | ✓ |
| ReadUntil via gRPC | ✗ | ✗ | ✓<sup>‡</sup> | ✓<sup>‡</sup> |
| Acceleration / parallelizable | ✗ / ✗ | (✓)<sup>†</sup> / ✓ | ✗ / ✓ | ✓ / ✓ |
| Open source | ✗ | ✓ | ✓ | ✓ |
| Fast generation of sequencing summary (no basecalling) | ✗ | ✗ | ✗ | ✓ |

Given the limitations of the existing simulators, the performance of SSDAs has been assessed by either evaluating the length of rejected reads and mapping speed as a proxy for selective sequencing performance (Firtina et al., 2023; Ulrich et al., 2022; Kovaka et al., 2021; Sneddon et al., 2022), or by running very few real experiments only (Kovaka et al., 2021; Payne et al., 2021). This is time-consuming, costs a non-negligible amount of money and prevents parameter optimization. Our simulator addresses this gap and allows to evaluate and optimize existing SSDAs more thoroughly whilst being lightweight. It can extract parameters from an existing run to be close yet not too similar to avoid overfitting the parameters to a particular run.

## 2. Methods

We present *SimReadUntil* which takes as input a set of full reads that can come from an existing run or from a full read simulator like NanoSim (Yang et al., 2017), see supplementary Figure 2. The reads may include adapter and barcode sequences. The (shuffled) full reads are distributed to the channels and short and long gaps are inserted between reads, where a long gap signifies a temporarily inactive channel. A channel can also become permanently inactive, not delivering any new reads. Each read has a start delay to account for adapters and barcodes. Mux scans can be triggered which terminate the current element in-progress across all channels (and are useful for simulating ultra-long read breaking). If a long gap is active when the mux scan starts, the long gap continues after the mux scan finishes. The simulator supports the ReadUntil API (Loose et al., 2016). The function `get_basecalled_read_chunks(channel)` returns any new chunks of the read *in-progress* that have not yet been returned. Different from the ONT ReadUntil API, the chunks are returned in basecalled format, after an added delay to mimic the basecalling delay (proportional to the number of nucleotides). We also offer conversion to a raw signal using ONT’s pore models to work with raw signal-based SSDAs (https://github.com/nanoporetech/kmer_models). A read no longer returns its chunks after the *stop receiving* action has been called on it, yet the full read is still written to disk. The *unblock* (reject) action terminates a read and adds an extra unblock delay right after it. If the read is no longer present by the time the action is processed, the action is logged as missed and signaled through the return value of the action call. Whenever a batch of reads has finished, the reads are written as FASTA records to disk. The FASTA header includes information such as the original full read id, the channel and the reason the read was ended, which is useful for post-processing and for later read mapping. The FASTA header contains the metadata to construct a sequencing summary file with the essential columns. For rejected reads with a NanoSim read id, the id is modified to reflect the effective reference length. The simulation can be accelerated and issues a warning if it cannot keep up with the desired acceleration. The ReadUntil API is supplemented with a gRPC implementation which makes the simulator usable from a wide range of programming languages and allows decoupling the simulator from the SSDA through interprocess communication.

Each channel collects statistics such as the number of reads, basepairs, rejections, missed rejections. They are output at regular intervals. The gaps are sampled from a `GapSampler` that receives the channel statistics as input. We provide several gap samplers that determine their parameters from an existing run. One gap sampler imitates an existing run (similarly to UNCALLED), another produces constant or random gaps for prototyping, yet another aggregates gaps from all channels and then samples them in fixed windows. Our recommended gap sampler `rolling_window_per_channel` samples the channel first (from the original run) and then samples gaps from a rolling window. New gap samplers can be easily added. See the Supplemental Material for more details.

The simulator enables benchmarking and hyperparameter tuning of selective sequencing algorithms. The hyperparameters can be tuned to different ONT devices, e.g., a GridION with a GPU can compute more than a portable MinION/Flongle that relies on a computer. Different policies for changing hyperparameters over the course of a run can also be tested. Moreover, new actions, such as stopping to receive data for a limited amount of time only, can be tested as a proof of concept for what real ONT devices should support.

## 3. Results

As opposed to MinKNOW and Icarust, our simulator is light-weight, so we run it on 2 cores of an Intel(R) Xeon(R) Gold 5220 CPU @ 2.20GHz with 8GB of RAM and 50GB of SSD disk. We ran one experiment comparing the different gap samplers and another that shows how to combine our simulator with ReadFish. We find that the `constant_gaps` method is a good starting point, but not very realistic (supplementary Figures 5, 6, 7). The `gap_replication` is the best method, but almost exactly reproduces a run, so SSDAs may memorize this structure and perform worse on a new run. Therefore, the recommended `rolling_window_per_channel` method represents a good trade-off between giving similar throughput and showing varying behavior at a smaller timescale. Similar to (Munro et al., 2023; Payne et al., 2021), we reproduce the enrichment of chromosome 21 and 22 with ReadFish connected to our simulator, as shown in supplementary Figures 8, 9. We demonstrate how to speed up the experiment by 10, replacing minimap2 alignment (Li, 2018) with ground-truth alignment.

## 4. Conclusion

We presented *SimReadUntil* to simulate an ONT device with support for the ReadUntil API, accessible both directly and via gRPC from a wide range of programming languages. The simulator only needs FASTA files of reads, which are available online and much smaller than bulk files or multi-fast5 files. Unlike other approaches which require often unavailable bulk or raw fast5 files, its parameters can be tuned based on a (comparatively small) sequencing summary file. By avoiding the basecalling step, the simulator allows to focus on the SSDA itself and removes the need for a GPU required by modern basecallers. This is beneficial when tuning approaches like ReadFish and ReadBouncer which operate directly on basecalled data.

We plan to use the simulator to establish a benchmark repository, optimize the hyperparameters of tools like ReadFish, and compare the performance on a real run with respect to default parameters. We are confident that our simulator can reliably predict the real-world performance of an SSDA in a new experiment. The modular design allows future work to come up with better gap samplers as needed, e.g., to model pore deterioration caused by rejections.

## Acknowledgements

We thank Bernhard Schölkopf for discussions.

We first compare existing simulators, then provide a schematic of our simulator, describe the approaches for parameter extraction from an existing sequencing summary file and compare the different parameter extraction methods, and finally present a use case that combines our simulator with ReadFish (and optionally NanoSim).

## A. Comparison of Existing Simulators

Table 2 compares existing simulators with ReadUntil capability. ONT’s MinKNOW controller software ships with a for-research simulator that takes a bulk file as input and plays it back. Read rejection is not very realistic: it simply cuts the read into two reads rather than taking a new read (Munro et al., 2023). Documentation is sparse and it requires non-trivial manual setup (Ulrich et al., 2022). The simulator from UNCALLED (Kovaka et al., 2021) is the approach closest to ours and comes with a Python wrapper around its C++ code. However, it is not well documented and not tested, does not write the finished reads to disk, often segfaults and is tightly integrated into their code, making it difficult to change simulation behavior beyond the obvious. It requires a big raw fast5 file (of several tens of GBs) and a sequencing summary file from an existing UNCALLED run, which is often unavailable. UNCALLED exactly replicates the channel activity pattern of active and inactive periods for a given channel and recycles reads which effectively means that no novel data is acquired since a selective sequencing run usually has many more reads than a normal run due to rejections. Icarust was developed in the same spirit as our simulator, but has drawbacks outlined in Table 2, mostly related to computational overhead associated with raw signals and no automatic (benchmarking-like) workflow to evaluate SSDA performance. Our simulator written in Python is probably also more accessible to modification than Icarust written in Rust.

**Table 2.**
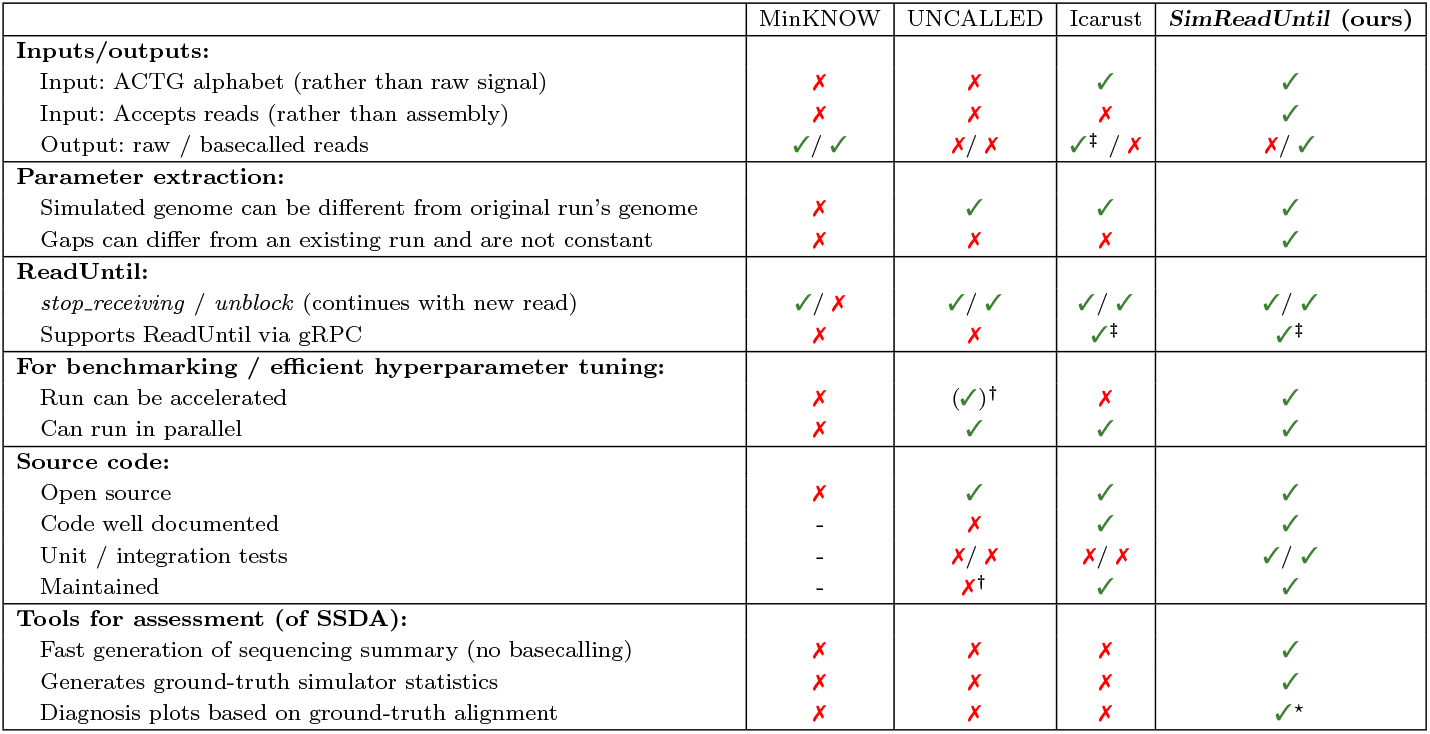
Comparison of simulators with ReadUntil functionality. Simulators without any ReadUntil functionality such as squiggle simulators are not shown. UNCALLED refers to the simulator of UNCALLED. Ground-truth simulator statistics include the number of rejected basepairs. Icarust was not yet officially published, so it may still change. *†*: UNCALLED’s acceleration mode distorts the length of long gaps. Icarust should be used instead of the UNCALLED simulator (personal communication). *‡*: The basecalled signal is converted to a raw signal using a pore model (or additionally with the memory-intensive Scrappie for Icarust). ⋆: See Figures 8, 9 for a subset of generated plots as well as the supplemental files.

Figure 1 gives an overview of the inputs and outputs for the existing tools. With minor modification, our simulator could also take raw signals as input. Due to the high raw signal sampling rate of 4kHz per channel, this requires a significant amount of memory, disk I/O and CPU processing (for fast5 extraction), which is relatively slow.^1^ This is unnecessary when benchmarking approaches that take basecalled reads like ReadFish and ReadBouncer, where we can instead simulate the sequencing errors in advance (e.g. with NanoSim) as we demonstrate in one of the use cases.

**Fig. 1:**
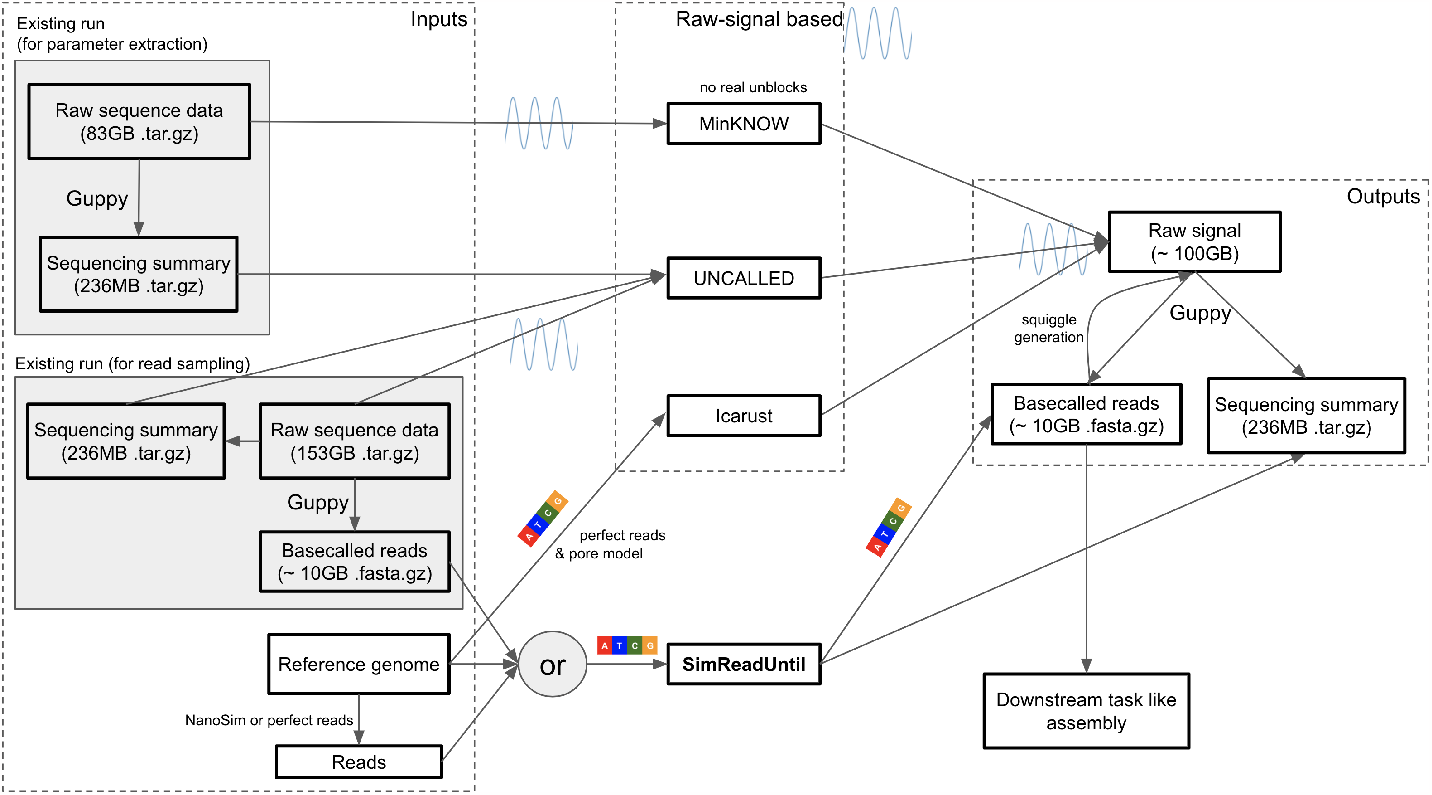
Inputs and outputs for the ReadUntil simulators. *SimReadUntil* and Icarust can take a reference genome and create perfect reads from them (with no mutations and indels) which are then input to the simulator. *SimReadUntil* also takes basecalled reads as input, which can be coming from an existing run or generated from a reference genome using NanoSim. UNCALLED takes both (full) raw reads and a sequencing summary (to locate the signal in the raw files) as input. The provided file sizes are taken from the UNCALLED HMW experiment (flowcell run 1) (Kovaka et al., 2021). The original run is 153GB in size whereas the selective sequencing run to mimic is 83GB in size. It has smaller file size because the throughput during selective sequencing is reduced due to more and longer gaps. Numbers prefixed with ∼ are estimated.

## B. Simulator Schematic

Figure 2 illustrates the design of the simulator as a collection of channels takes full reads and outputting (full or partial) reads according to ReadUntil actions. An example of the simulator with two channels with random ReadUntil actions is given in Figure 3.

**Fig. 2:**
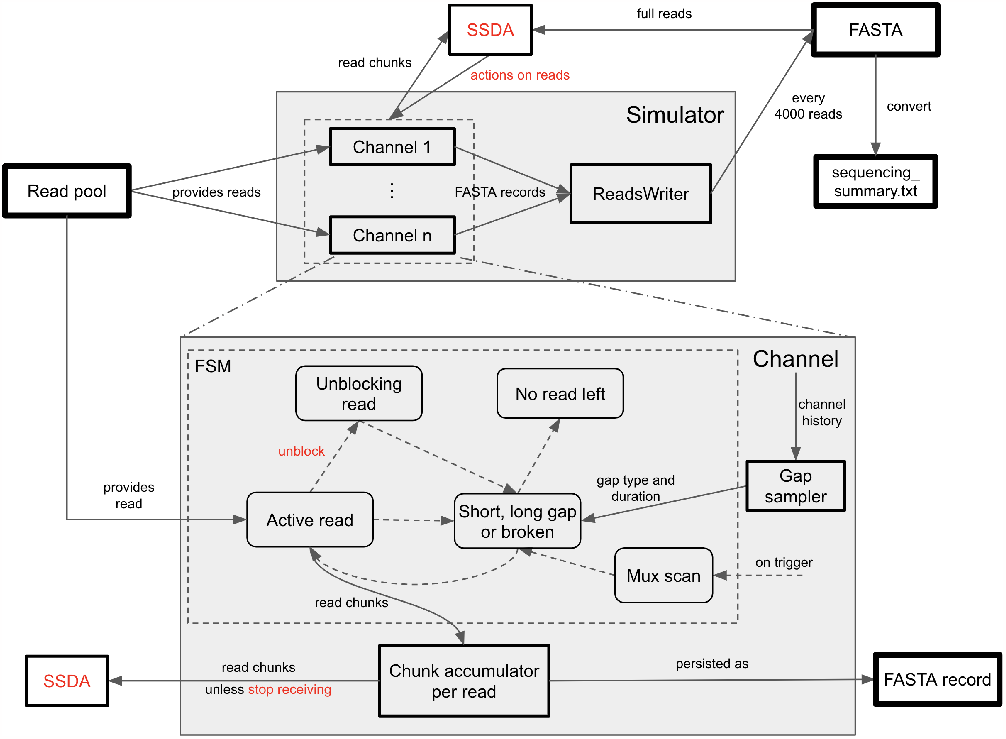
The top shows *SimReadUntil* which consists of a set of channels, the bottom zooms into a single channel. Solid arrows mean data is flowing. Inputs and outputs are shown in a square box with thick border. **Top:** Simulator grouping a set of channels that fetch reads from a reads pool, e.g., a reads file generated with NanoSim and possibly shuffled. Finished reads are output as a FASTA file every 4000 reads. The FASTA output can be converted into a sequencing summary file (which is commonly output by ONT basecallers) since the FASTA header contains all relevant information. **Bottom:** Zoom into a single channel which is a finite-state machine (FSM) whose transitions (dashed arrows) between states (rounded boxes) happen either due to time or ReadUntil decisions (red) made by the selective sequencing decision algorithm (SSDA). The gaps are chosen from a gap sampler and can depend on the channel history with parameters tuned from an existing run. The mux scan can be triggered at any time (ejecting the element in-progress) and is followed by a gap once it finishes.

**Fig. 3:**
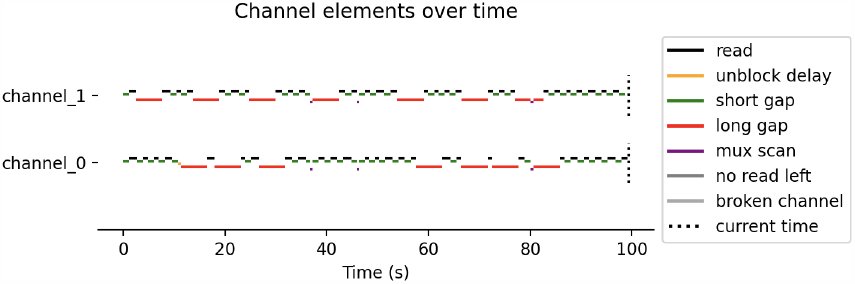
Example of a simulation with two channels. Parameters were chosen to show all available channel elements. In real ONT runs, mux scans occur roughly every 90 minutes.

ONT’s ReadUntil gRPC implementation `get_live_reads` streams read chunks as a bidirectional gRPC stream, even when the data is not requested. We instead provide the data upon request, thereby reducing computational overhead on the simulator if data is never requested. This assumes that the gRPC connection between the simulator and the SSDA under test is fast, which is the case, especially since the basecalled data is much smaller than raw signal data. If using Python, it is possible to directly connect to the simulator without the gRPC overhead.

### B.1. NanoSim & NanoSim Read IDs

We have modified the NanoSim implementation to make it deterministic (in single-process mode), removed flanking regions and made small optimizations to speed it up, e.g., the unnecessary big read error profile is not written to disk. NanoSim is a submodule of our git repository. We use the pre-trained error models from NanoSim which are saved in pickle format. Due to API changes in numpy and scipy, we could not load them into the same environment as our simulator, so we decided to create a separate conda environment as suggested in their setup.

NanoSim read ids contain the ground-truth alignment information. When simulating a genome (as opposed to a metagenome) without chimeric reads, the ids are of the form:

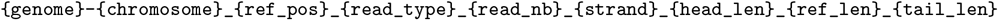

The read comes from the forward or reverse `strand` of `chromosome` in `genome` starting at 0-based position `ref_pos` with a length of `ref_len` on the forward strand of the reference, so it spans the half-open interval `[ref_pos:ref_pos+ref_len)` on the forward strand. When a read is rejected (*unblock*), the `ref_pos` and `ref_len` are adapted proportionally. NanoSim adds flanking regions around the read consisting of random letters. We instead introduced a read delay which stands for the adapter and poly A tail preceding the actual read, so we set `head_len` and `tail_len` to 0. The read number `read_nb` is unique to each read and concatenates the process id and read number (as NanoSim uses multiprocessing). The read type is `aligned` or `perfect` for reads with and without errors respectively.^2^ Since NanoSim is rather slow, we recommend pre-generating the reads (which can result in files on the order of 10GBs). It is possible to adapt the NanoSim implementation to output reads on the fly, provided enough cores are used to keep up with the simulation. (In the metagenomic setting, this requires some extra work due to process synchronization to ensure a metagenomic abundance ratio.)

Our simulator comes with post-processing tools that parse the read ids and plot the fraction of basepairs covered at least *x* times for various values of *x* over time per `{genome}-{chromosome}`. When the reference genome is large, we reduce the granularity during counting by taking blocks of basepairs for efficiency.

## C. Parameter Extraction Methods

We set the parameters of the simulator based on the `sequencing_summary.txt` of an existing run. The sequencing summary is available from an ONT run after basecalling the reads from the raw signal. This provides information of the timings of the reads in each channel. We extract the following parameters:

- The number of basepairs per second `bp_per_second` is estimated as the median read speed.
- The read delay is the delay between the start of a read until sequencing basepairs from the sample that are not part of the adapter or barcode, and can be computed as the difference between the template start and the read start. We set it to be the median over all observed delays.
- The unblocking delay is the duration during which the unblock voltage is applied to eject the read. This is typically set to a fixed value ≈ 0.1s (Kovaka et al., 2021).

We decided to set these parameters to constant values because they are mostly constant and to limit the number of parameters.

The gap sampler returns the length of the gap following a read as well as the gap type (long or short). Similarly to UNCALLED, we classify the gaps as short and long gaps. The threshold to determine this is the median plus five times the interquartile range of the gap lengths computed over all observed gaps across all channels.^3^ A long gap continues after a mux scan, whereas a short gap does not. Gaps are the regions between reads in the interval 0 to the end time of the last read of a sequencing run. We have implemented the following gap samplers whose parameters are fitted from an existing run:

- `random_gaps` samples gaps from an arbitrarily chosen (fixed) distribution and does not extract parameters from an existing run. This is for debugging and prototyping new methods.
- `constant_gaps` computes the median of the short gaps and long gaps respectively, and selects a long gap with a constant probability. Due to the skewed gap length distribution (towards long gap durations), setting the probability of a long gap to be equal to the fraction of long gaps is not appropriate because this means that the fraction of time spent in short gaps becomes larger. Rather, we require the fraction of time spent in long gaps to be the same as in the original run. For this to be the case, we set the probability of selecting a long gap to be 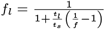, where *t*_*s*_ and *t*_*l*_ are the times of short and long gaps respectively and *f* is the fraction of long gaps among all gaps. We provide a derivation below. This approach is useful for rapid prototyping. It is similar to Icarust which employs a constant (configurable) short gap defaulting to 0.4s. Unlike Icarust, we extract this delay from the data and distinguish short from long gaps.
- `gap_replication` replicates gaps similarly to UNCALLED. A channel consists of alternating active and inactive periods. The long gaps represent long inactive periods without any reads, the time between them corresponds to the active periods. The order and duration of active and inactive periods is replicated as in the real run. Within an active period, gaps between reads are sampled from the short gaps observed within that period. A period ends once its duration is exceeded. This waits until the read has finished, so an active period may be longer than in a real run. It is suited when we want to replicate an existing run as well as possible, but still allow for ReadUntil decisions to influence the run. This method is similar to UNCALLED and MinKNOW (which does not support unblocks properly).
- `window_all_channels` defines contiguous time windows and samples the gaps from the corresponding time windows in the real run. To avoid abrupt changes, the gaps are sampled from overlapping windows, see Figure 4. The windowed approach ensures that gaps from the start and end of a sequencing run are not mixed because they have different statistics: gaps at the end are considerably longer because the pores deteriorate. The gaps are sampled from the data in the time window over all channels to have sufficient data, but this mixes good and bad channels.
- **Recommended method**: `rolling_window_per_channel` operates per channel and uses rolling windows compared to `window_all_channels`. It first samples the channel in the original run, then samples gaps in a rolling time window around the current timepoint in the corresponding channel, see Figure 4. It stops once the sequencing time of the channel is exceeded. The disadvantage of sampling gaps per channel is that there are fewer gaps in the window (as low as 1) especially towards the end when the channel is more likely to be inactive for long periods of time.

**Fig. 4:**
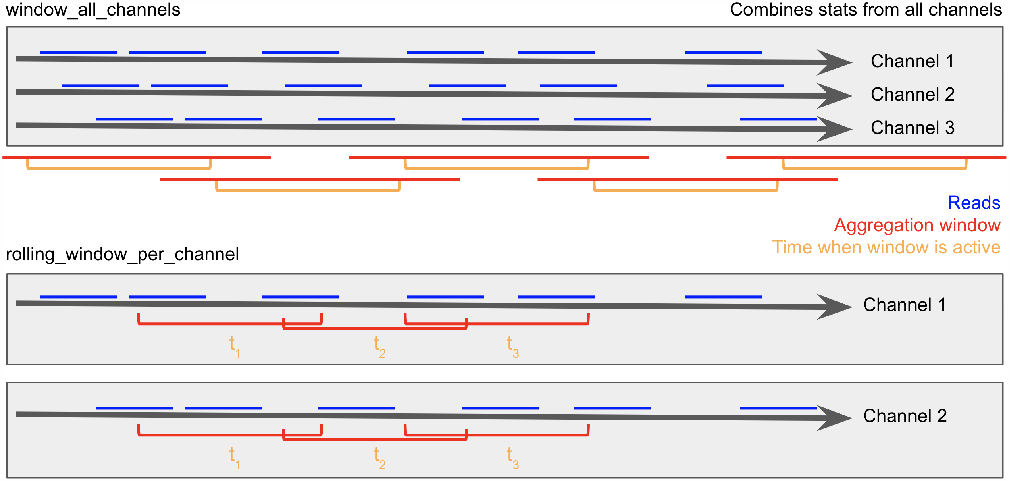
Gap sampler methods extracting gaps between reads. Top: `window_all_channels` gap sampler with contiguous time windows that sample gaps from the aggregation windows containing them, so the aggregation windows between adjacent time windows overlap. The windows aggregate data over all channels, so well-performing and badly-performing channels are mixed. Bottom: `rolling_window_per_channel` gap sampler with rolling windows per channel (only showing 2 channels for simplicity). The per-channel sampling means less data is available, especially towards the end of the sequencing run.

The number of channels can be flexibly chosen except for `gap_replication`. Since the `gap_replication` approach is exactly replicating an existing run, an SSDA may overfit to the existing run. Channels deteriorate over time and some channels work better than others. To take this into account, all methods except `random_gaps` sample from the empirical distribution of sequencing time per channel. Once the time is exceeded (after a read has finished), the channel stops producing any new reads. We also experimented with fitting the time until a channel produces its last read to a distribution (beta, weibull min), but this did not deliver equally good results in terms of basepairs and reads over the full run. Since loading the sequencing summary for parameter extraction takes some time (though *<* 1 minute on a normal computer for a 200-500MB sized sequencing summary), we cache the simulation parameters.

We now derive the probability of sampling from a long gap to ensure that the fraction of time spent in long gaps corresponds to the fraction 0 ≤ *f* ≤ 1 of an existing run. Assume we have sampled *n* gaps with 0 ≤ *x*_*l*_ ≤ *n* long gaps among them. Let *t*_*s*_, *t*_*l*_ denote the expected length of short and long gaps respectively^4^. We want the following to hold in expectation:

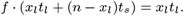

Note that we do not match the ratio, but multiply the denominator because the expectation of a ratio cannot be easily simplified with linearity (*x*_*l*_ is a random variable). Taking the expectation, we have 𝔼 [*x*_*l*_] = *f*_*l*_*n* given that we independently sample a long gap with probability *f*_*l*_. Inserting this, we solve for *f*_*l*_:

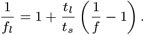

As a sanity check, the case *t*_*s*_ = *t*_*l*_ corresponds to *f*_*l*_ = *f, t*_*l*_*/t*_*s*_ → ∞ to *f*_*l*_ → 0, *t*_*l*_*/t*_*s*_ → 0 to *f*_*l*_ → 1, *f* → 0 to *f*_*l*_ → 0, *f* → 1 to *f*_*l*_ → 1.

### C.1. Comparison of Parameter Extraction Methods from a Real Run

In this section, we extract parameters from a real run and compare the statistics of the simulated run to the original run, see `replicate_run.py`. For this, we take the sequencing summary file provided by UNCALLED (Kovaka et al., 2021) mentioned in the README of their GitHub: https://labshare.cshl.edu/shares/schatzlab/www-data/UNCALLED/simulator_files/20190809_zymo_seqsum.txt.gz from a sequencing run of the Zymo mock community. We run the simulated run without selective sequencing since selective sequencing affects the statistics like read duration. Since mux scans potentially introduce long gaps that we do not want to model, we first find and remove mux scans with a script that we have adapted from UNCALLED.^5^ We then extract the read duration from the sequencing summary file and generate reads such that their lengths match the extracted read durations (at 450 bp s^*−*1^ speed). The simulator forwards each channel in parallel. The implementation is parallelized with *joblib* and faster the more cores are available. Note that we only parallelized the code when ReadUntil functionality is not used, which is the case here.

We compare the different parameter extraction methods. More figures are available in the supplementary material. Figure 5 compares the number of actively reading channels at regular timepoints. We see that the `gap_replication` closely matches the original run `zymo_real_run`. The `rolling_window_per_channel` also works very well while offering some deviation from the existing run. The `window_all_channels` works well at the beginning, less at the end. A reason may be that the method averages across all channels, thus combining information from channels with different channel healths. The `constant_gaps` strategy does not perform well, especially at the end. Figure 6 compares the number of reads and sequenced basepairs over time. Figure 7 compares the channel statistics for the various methods. We see that the qualitative behavior is similar to before.

**Fig. 5:**
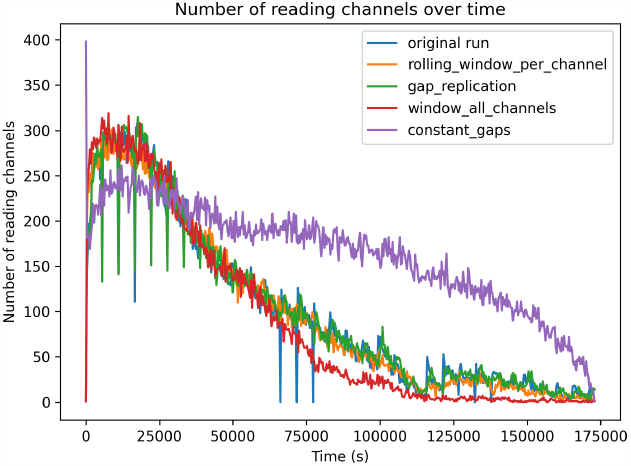
Actively reading channels over time. The original sequencing run is `zymo_real_run`.

**Fig. 6:**
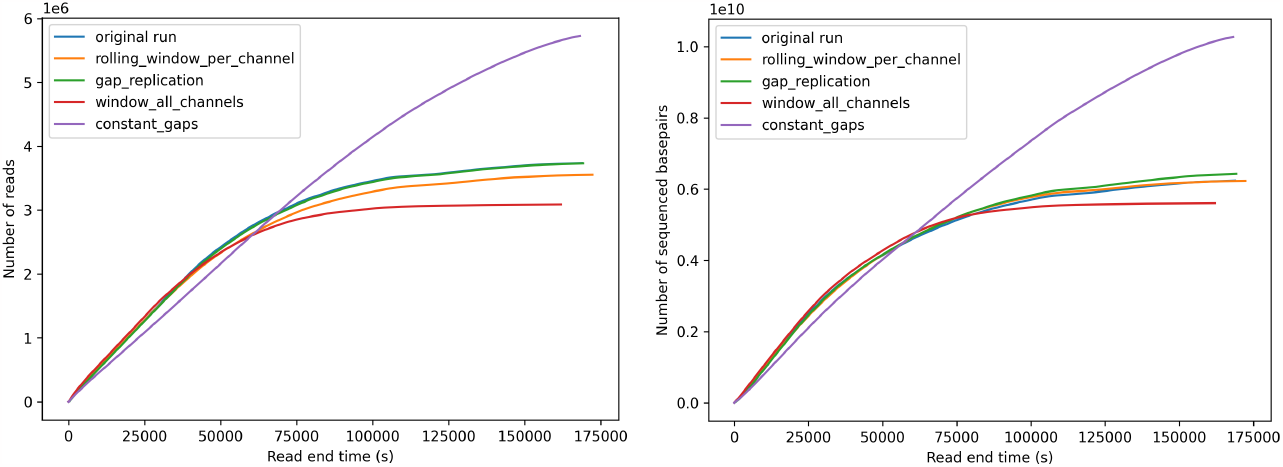
Total number of reads and total number of sequenced basepairs across all channels over time. The original sequencing run `zymo_real_run` and the replication run `gap_replication` are overlapping as expected. Some curves overlap.

**Fig. 7:**
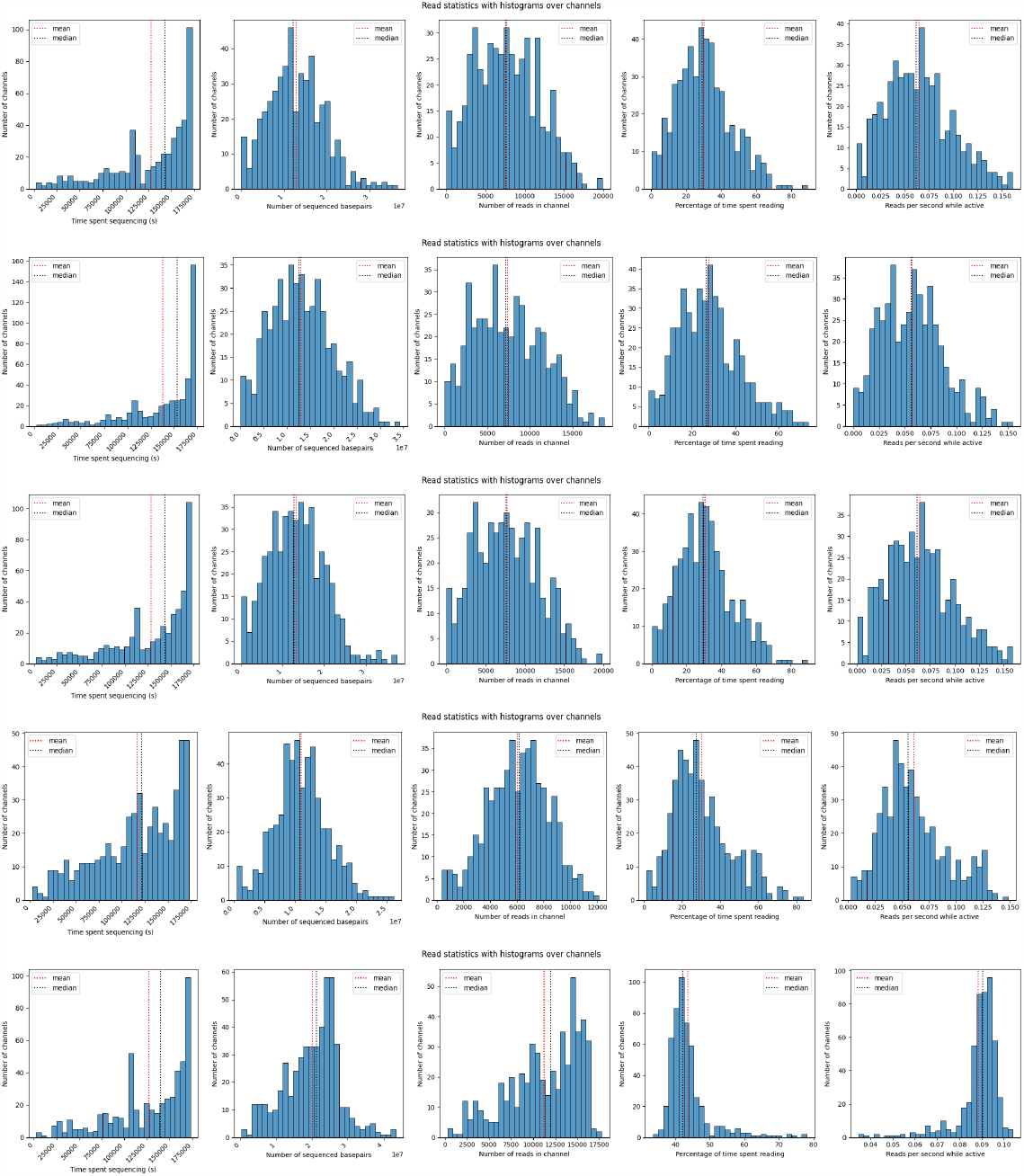
Statistics across channels for the `zymo_real_run` (original run), `rolling_window_per_channel` (recommended method), `gap_replication` (replication), `window_all_channels` (mixing all channels) and `constant_gaps` (constant gaps) parameter extraction methods (top to bottom). The percentage of time spent reading only accounts for the percentage until the channel produces its last read.

## D. Use Case: SimReadUntil with ReadFish

We present an example on how to use our simulator with ReadFish (Payne et al., 2021), see `enrich_usecase.py`. ReadFish takes selective sequencing decisions by mapping reads to a reference with *minimap2* (Li, 201*8). Similar to other work (Payne et al*., *2021;* Munro et al.,2023), we take the human reference genome and enrich chromosomes 20 and 21. To focus on the simulator and not be affected by mapping errors, we generate perfect reads on-the-fly by randomly choosing the chromosome, start location, length (between 5000 and 10000) and strand (forward/reverse). The simulation parameters are extracted from the same Zymo mock sequencing summary file as above using the `rolling_window_per_channel` parameter extraction method after removing mux scans. We simulate 512 channels with acceleration factor 10. We slightly modified ReadFish’s implementation to feed in extra parameters, without modifying its behavior. In particular, we provide a fake basecaller that returns the basecalled data as-is since our modified ReadUntil API returns basecalled data rather than raw signal data. Since basecalling constitutes a fundamental bottleneck of ReadFish, we introduce a time delay per basecalled basepair that we estimated from benchmarking experiments (Benton, 2023). A GPU can call roughly *2*.6 × 10^7^ samples s^*−*1^, where a sample refers to the ONT raw signal at 4 kHz producing roughly 450 bp s^*−*1^. This means (4 kHz*/*450 bp s^*−*1^)*/*(2.6 × 10^7^ samples s^*−*1^) = 3.4 × 10^*−*7^ s bp^*−*1^. In accelerated mode, we divide this number by the acceleration factor. In each iteration, ReadFish requests new read chunks from all 512 channels (batch size 512), basecalls them, maps them to the reference with `minimap2`, makes decisions about each of them, then communicates the decisions in one batch via the ReadUntil API and waits to ensure a minimum time between iterations (throttle). In accelerated mode, we observed that the batch size is too big in the sense that the latency (but not throughput) is too high, making selective sequencing decisions arrive too late. This illustrates how the simulator can be used to find a good batch size. By the time the read rejections are communicated, 75% of the reads have already finished, which greatly reduces the effectiveness of ReadFish. Therefore, we compute the batch size as the number of channels divided by the acceleration factor. We divide the default throttle of 0.1 s by the square of the acceleration factor (roughly).^6^ We noticed that *minimap2* is a bottleneck at acceleration factor 10. Therefore, we replace *minimap2* by a fake aligner that parses the NanoSim read id to find the mapping position. We do not add any mapping delay (taking into account the acceleration factor) as of now.

From Figures 8, 9, we observe that ReadFish is effective at enriching chromosomes 20 and 21 in this ideal setting of perfect reads and low-latency alignment. Chromosomes 20 and 21 (named “target”) are enriched whereas reads coming from other chromosomes are rejected, mostly after 500 bps. Without selective sequencing, the ratio of target to non-target basepairs would be 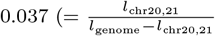 with *l* being the length), whereas here, we observe roughly 0.58 estimated from the plots, which is a 16-fold improvement. This is very optimistic because we assume perfect basecalling and mapping. The fraction covered is computed by dividing each chromosome into blocks of size `blocksize` (except for the last which is shorter). Whenever a read intersects with a block, we increment that block’s count by the number of basepairs intersecting with the read. After adding all reads up to time t, we can compute the fraction of blocks with average coverage at least 1, 2, 3, 4 which is shown in the figure comparing the target to the non-target chromosomes. We use a `blocksize` of 1000 which greatly speeds up computations with respect to `blocksize` 1.

**Fig. 8:**
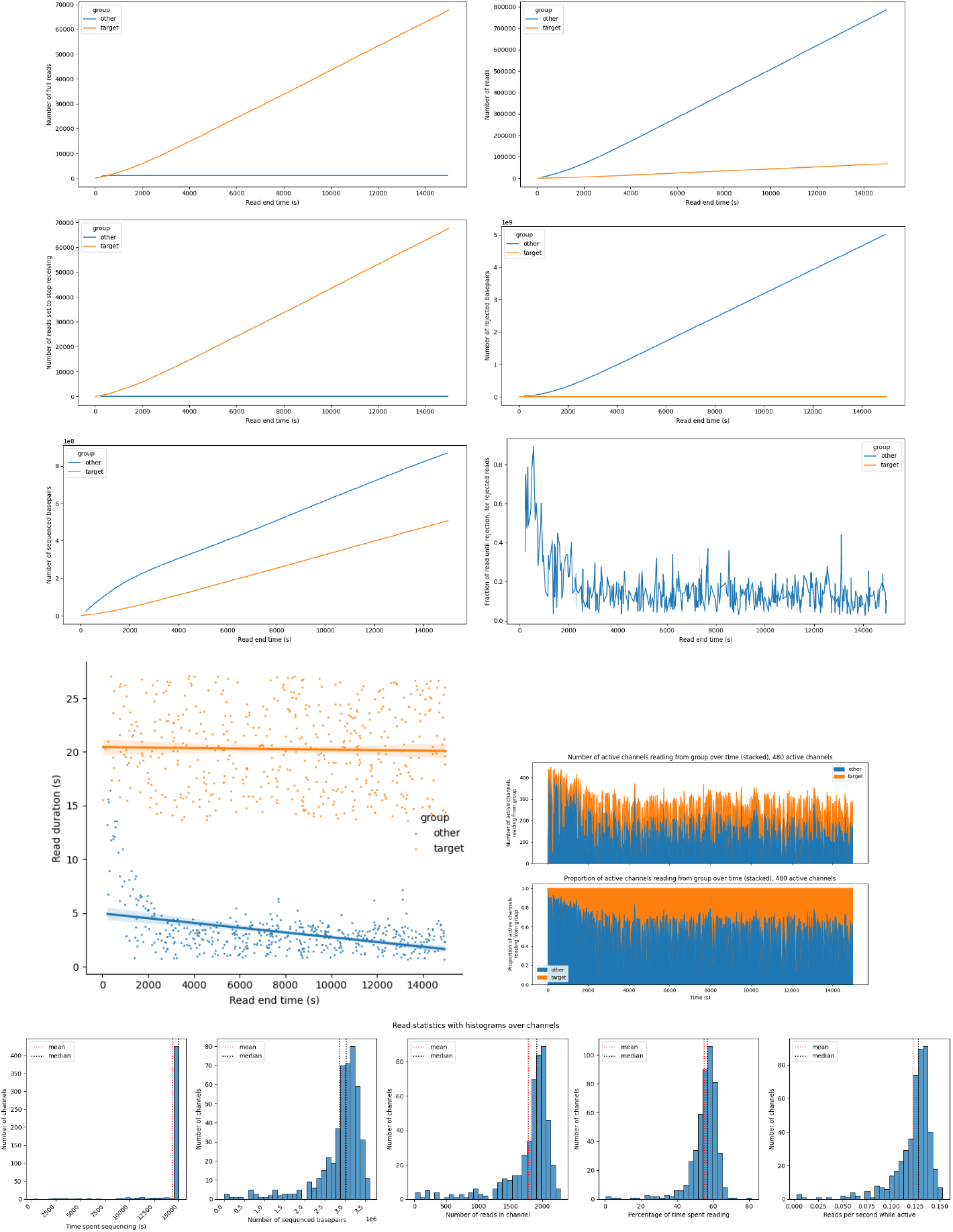
Combining the simulator with ReadFish to enrich for chromosomes chr20, 21 (target vs other). Number of reads, full reads (reads that were not rejected), reads set to *stop receiving*, number of rejected and sequenced basepairs, fraction of read until it got rejected, read duration of reads, number and relative proportion of channels reading from target vs other, read stats by channel. The simulation is running for 15000s which is significantly less than the flowcell lifetime, so almost all channels sequence until the end. The read end time of the target group does not start at zero because the reads from the target group do not get rejected, whereas the non-target reads get rejected immediately.

**Fig. 9:**
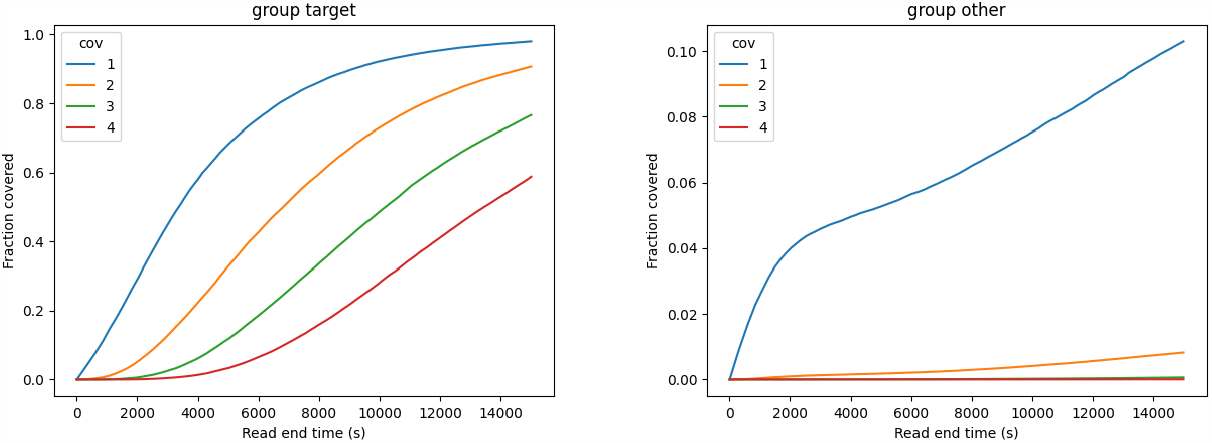
Combining the simulator with ReadFish to enrich for chromosomes chr20, 21 (target vs other). Fraction of basepairs covered at least *x* times for *x* = 1, 2, 3, 4 as the sequencing run progresses. Coverage for target is much better than for the rest.

Our script also permits to generate reads with NanoSim and feed them to the simulator. We do not do it here since NanoSim read generation is rather slow. We provide a jupyter script that shows how to generate reads with NanoSim. Note that we fixed bugs in NanoSim to make runs with the same random seed reproducible and made other small modifications, available as a submodule in our repository, see Section B.1.

## E. Extra Files

The supplemental material contains additional figures from the experiments described here.

## Footnotes

1 UNCALLED (Kovaka et al., 2021) mentions that their simulator can take 100GB of memory, see https://github.com/skovaka/UNCALLED#readme.

2 For perfect reads, it is easier and more efficient to generate the reads directly in-memory as input to the simulator.

3 This is different from UNCALLED which uses the median plus one standard deviation. Since the gap length distribution has many outliers, the standard deviation is not a very robust measure.

4 Here, we derive it for the general case when the gap length is not necessarily constant, e.g., sampled from some distribution. In the setting above, we extract the medians, so the short and long gap times are constant.

5 This script is not perfect, which explains jumps in the plots of actively reading channels around the mux scan intervals. The mux scans occur roughly 90 minutes, but not exactly.

6 We divide by the square to account for both the acceleration and the reduced batch size.

