## Supplement for "SimReadUntil for Benchmarking Selective Sequencing Algorithms on ONT Devices"

| | MinKNOW | UNCALLED | Icarust | ***SimReadUntil* (ours)** |
|---|---|---|---|---|
| **Inputs/outputs:** | | | | |
| Input: ACTG alphabet (rather than raw signal) | ✗ | ✗ | ✓ | ✓ |
| Input: Accepts reads (rather than assembly) | ✗ | ✗ | ✗ | ✓ |
| Output: raw / basecalled reads | ✓/ ✓ | ✗/ ✗ | ✓‡ / ✗ | ✗/ ✓ |
| **Parameter extraction:** | | | | |
| Simulated genome can be different from original run's genome | ✗ | ✓ | ✓ | ✓ |
| Gaps can differ from an existing run and are not constant | ✗ | ✗ | ✗ | ✓ |
| **ReadUntil:** | | | | |
| *stop_receiving* / *unblock* (continues with new read) | ✓/ ✗ | ✓/ ✓ | ✓/ ✓ | ✓/ ✓ |
| Supports ReadUntil via gRPC | ✗ | ✗ | ✓‡ | ✓‡ |
| **For benchmarking / efficient hyperparameter tuning:** | | | | |
| Run can be accelerated | ✗ | (✓)† | ✗ | ✓ |
| Can run in parallel | ✗ | ✓ | ✓ | ✓ |
| **Source code:** | | | | |
| Open source | ✗ | ✓ | ✓ | ✓ |
| Code well documented | - | ✗ | ✓ | ✓ |
| Unit / integration tests | - | ✗/ ✗ | ✗/ ✗ | ✓/ ✓ |
| Maintained | - | ✗† | ✓ | ✓ |
| **Tools for assessment (of SSDA):** | | | | |
| Fast generation of sequencing summary (no basecalling) | ✗ | ✗ | ✗ | ✓ |
| Generates ground-truth simulator statistics | ✗ | ✗ | ✗ | ✓ |
| Diagnosis plots based on ground-truth alignment | ✗ | ✗ | ✗ | ✓⋆ |

**Table 1.** Comparison of simulators with ReadUntil functionality. Simulators without any ReadUntil functionality such as squiggle simulators are not shown. UNCALLED refers to the simulator of UNCALLED. Ground-truth simulator statistics include the number of rejected basepairs. Icarust was not yet officially published, so it may still change. †: UNCALLED's acceleration mode distorts the length of long gaps. Icarust should be used instead of the UNCALLED simulator (personal communication). ‡: The basecalled signal is converted to a raw signal using a pore model (or additionally with the memory-intensive Scrappie for Icarust). ⋆: See Figures 8, 9 for a subset of generated plots as well as the supplemental files.

[1] UNCALLED (Kovaka et al., 2021) mentions that their simulator can take 100GB of memory, see `https://github.com/skovaka/UNCALLED#readme`.

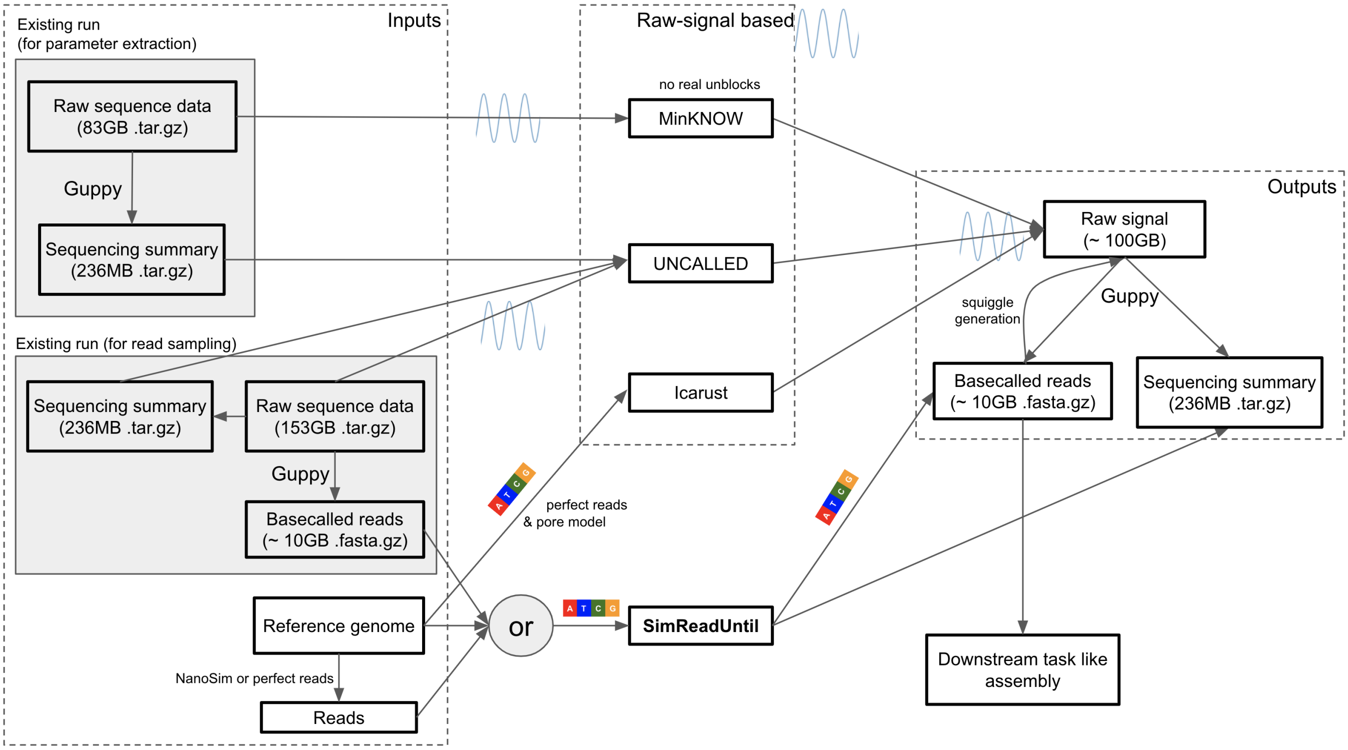


Fig. 1: Inputs and outputs for the ReadUntil simulators. *SimReadUntil* and Icarust can take a reference genome and create perfect reads from them (with no mutations and indels) which are then input to the simulator. *SimReadUntil* also takes basecalled reads as input, which can be coming from an existing run or generated from a reference genome using NanoSim. UNCALLED takes both (full) raw reads and a sequencing summary (to locate the signal in the raw files) as input. The provided file sizes are taken from the UNCALLED HMW experiment (flowcell run 1) (Kovaka et al., 2021). The original run is 153GB in size whereas the selective sequencing run to mimic is 83GB in size. It has smaller file size because the throughput during selective sequencing is reduced due to more and longer gaps. Numbers prefixed with ∼ are estimated.

ONT's ReadUntil gRPC implementation `get_live_reads` streams read chunks as a bidirectional gRPC stream, even when the data is not requested. We instead provide the data upon request, thereby reducing computational overhead on the simulator if data is never requested. This assumes that the gRPC connection between the simulator and the SSDA under test is fast, which is the case, especially since the basecalled data is much smaller than raw signal data. If using Python, it is possible to directly connect to the simulator without the gRPC overhead.

### B.1. NanoSim & NanoSim Read IDs

We have modified the NanoSim implementation to make it deterministic (in single-process mode), removed flanking regions and made small optimizations to speed it up, e.g., the unnecessary big read error profile is not written to disk. NanoSim is a submodule of our git repository. We use the pre-trained error models from NanoSim which are saved in pickle format. Due to API changes in numpy and scipy, we could not load them into the same environment as our simulator, so we decided to create a separate conda environment as suggested in their setup.

NanoSim read ids contain the ground-truth alignment information. When simulating a genome (as opposed to a metagenome) without chimeric reads, the ids are of the form:

```
{genome}-{chromosome}_{ref_pos}_{read_type}_{read_nb}_{strand}_{head_len}_{ref_len}_{tail_len}
```

The read comes from the forward or reverse `strand` of `chromosome` in `genome` starting at 0-based position `ref_pos` with a length of `ref_len` on the forward strand of the reference, so it spans the half-open interval `[ref_pos:ref_pos+ref_len)` on the forward strand. When a read is rejected (*unblock*), the `ref_pos` and `ref_len` are adapted proportionally. NanoSim adds flanking regions around the read consisting of random letters. We instead introduced a read delay which stands for the adapter and poly A tail preceding the actual read, so we set `head_len` and `tail_len` to 0. The read number `read_nb` is unique to each read and concatenates the process id and read number (as NanoSim uses multiprocessing). The read type is `aligned` or `perfect` for reads with and without errors respectively.[2] Since NanoSim is

[2] For perfect reads, it is easier and more efficient to generate the reads directly in-memory as input to the simulator.

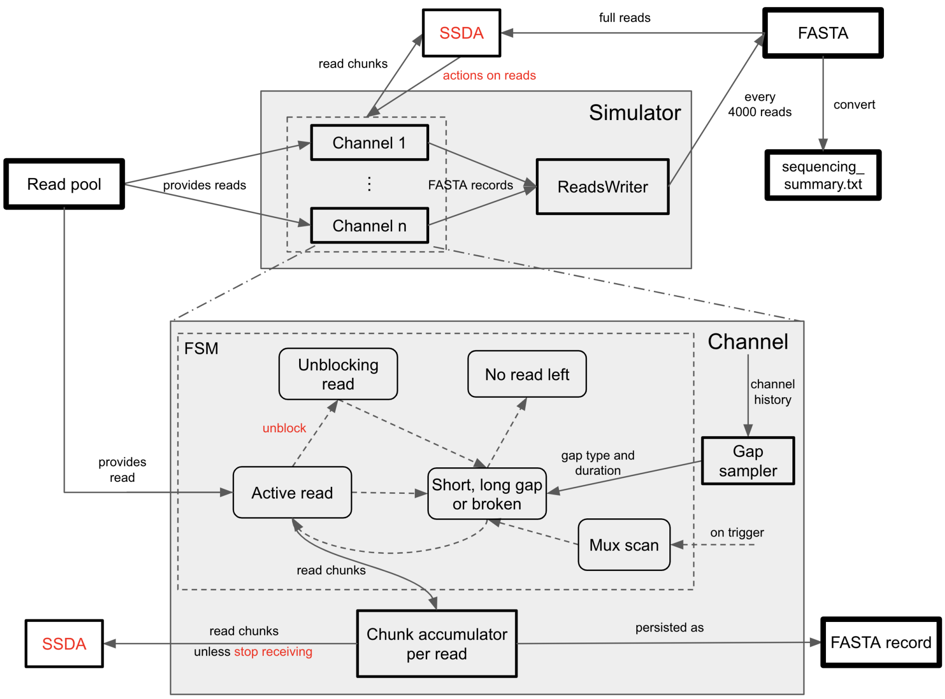


Fig. 2: The top shows *SimReadUntil* which consists of a set of channels, the bottom zooms into a single channel. Solid arrows mean data is flowing. Inputs and outputs are shown in a square box with thick border. **Top:** Simulator grouping a set of channels that fetch reads from a reads pool, e.g., a reads file generated with NanoSim and possibly shuffled. Finished reads are output as a FASTA file every 4000 reads. The FASTA output can be converted into a sequencing summary file (which is commonly output by ONT basecallers) since the FASTA header contains all relevant information. **Bottom:** Zoom into a single channel which is a finite-state machine (FSM) whose transitions (dashed arrows) between states (rounded boxes) happen either due to time or ReadUntil decisions (red) made by the selective sequencing decision algorithm (SSDA). The gaps are chosen from a gap sampler and can depend on the channel history with parameters tuned from an existing run. The mux scan can be triggered at any time (ejecting the element in-progress) and is followed by a gap once it finishes.

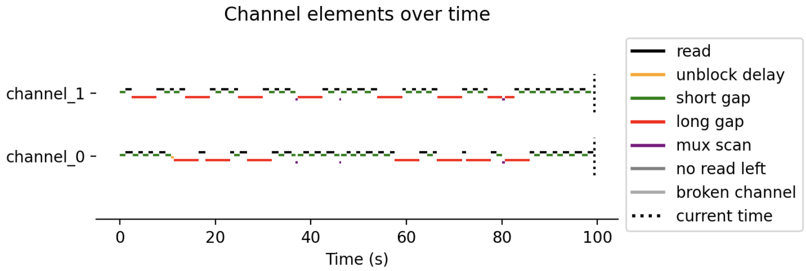


Fig. 3: Example of a simulation with two channels. Parameters were chosen to show all available channel elements. In real ONT runs, mux scans occur roughly every 90 minutes.

rather slow, we recommend pre-generating the reads (which can result in files on the order of 10GBs). It is possible to adapt the NanoSim implementation to output reads on the fly, provided enough cores are used to keep up with the simulation. (In the metagenomic setting, this requires some extra work due to process synchronization to ensure a metagenomic abundance ratio.)

We now derive the probability of sampling from a long gap to ensure that the fraction of time spent in long gaps corresponds to the fraction $0 \leq f \leq 1$ of an existing run. Assume we have sampled $n$ gaps with $0 \leq x_l \leq n$ long gaps among them. Let $t_s, t_l$ denote the

[3] This is different from UNCALLED which uses the median plus one standard deviation. Since the gap length distribution has many outliers, the standard deviation is not a very robust measure.

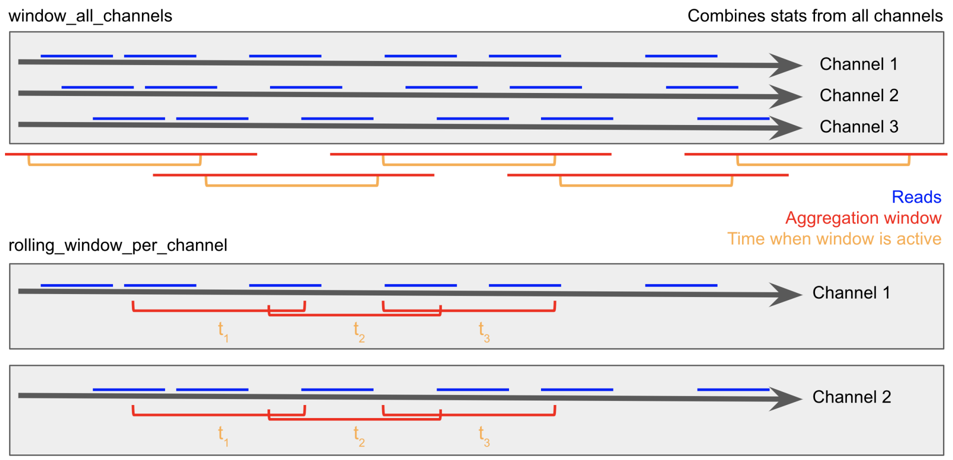


Fig. 4: Gap sampler methods extracting gaps between reads. Top: `window_all_channels` gap sampler with contiguous time windows that sample gaps from the aggregation windows containing them, so the aggregation windows between adjacent time windows overlap. The windows aggregate data over all channels, so well-performing and badly-performing channels are mixed. Bottom: `rolling_window_per_channel` gap sampler with rolling windows per channel (only showing 2 channels for simplicity). The per-channel sampling means less data is available, especially towards the end of the sequencing run.

expected length of short and long gaps respectively[4]. We want the following to hold in expectation:

$$f \cdot (x_l t_l + (n - x_l) t_s) = x_l t_l.$$

Note that we do not match the ratio, but multiply the denominator because the expectation of a ratio cannot be easily simplified with linearity ($x_l$ is a random variable). Taking the expectation, we have $\mathbb{E}[x_l] = f_l n$ given that we independently sample a long gap with probability $f_l$. Inserting this, we solve for $f_l$:

$$\frac{1}{f_l} = 1 + \frac{t_l}{t_s}\left(\frac{1}{f} - 1\right).$$

As a sanity check, the case $t_s = t_l$ corresponds to $f_l = f$, $t_l/t_s \to \infty$ to $f_l \to 0$, $t_l/t_s \to 0$ to $f_l \to 1$, $f \to 0$ to $f_l \to 0$, $f \to 1$ to $f_l \to 1$.

#### C.1. Comparison of Parameter Extraction Methods from a Real Run

[5] This script is not perfect, which explains jumps in the plots of actively reading channels around the mux scan intervals. The mux scans occur roughly 90 minutes, but not exactly.

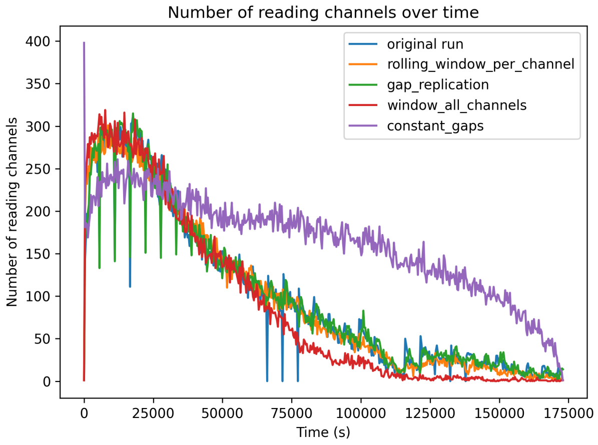


Fig. 5: Actively reading channels over time. The original sequencing run is **zymo_real_run** .

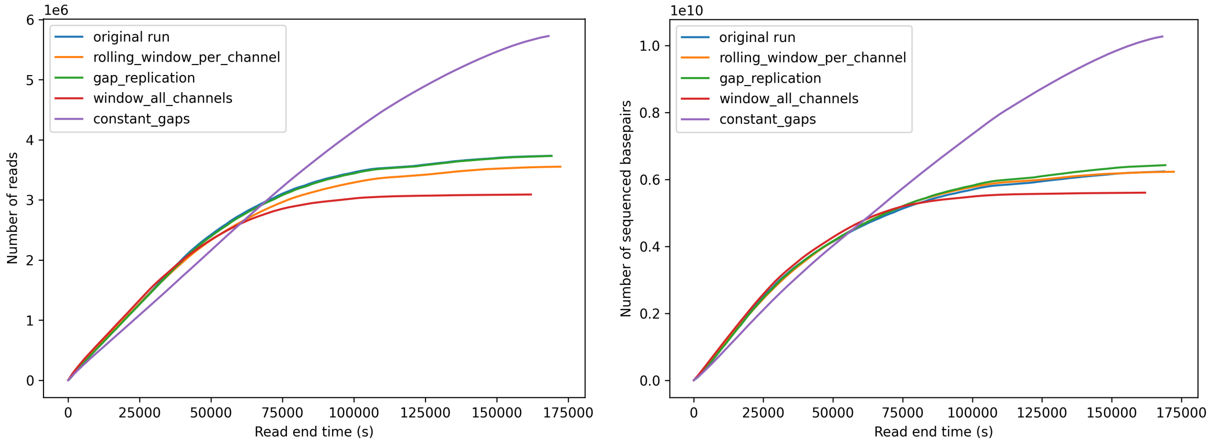


Fig. 6: Total number of reads and total number of sequenced basepairs across all channels over time. The original sequencing run **zymo_real_run** and the replication run **gap_replication** are overlapping as expected. Some curves overlap.

[6] We divide by the square to account for both the acceleration and the reduced batch size.

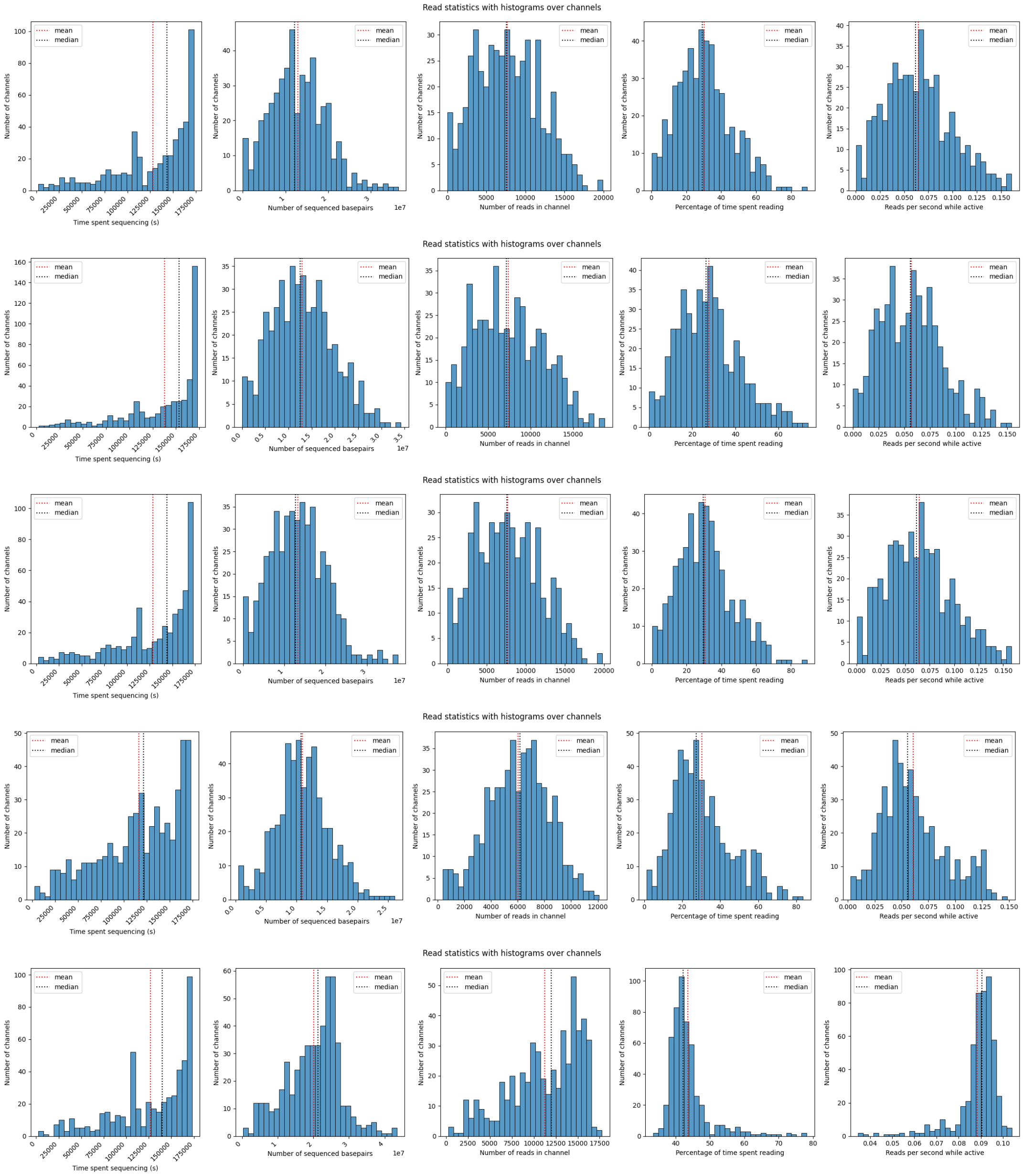


Fig. 7: Statistics across channels for the **zymo_real_run** (original run), **rolling_window_per_channel** (recommended method), **gap_replication** (replication), **window_all_channels** (mixing all channels) and **constant_gaps** (constant gaps) parameter extraction methods (top to bottom). The percentage of time spent reading only accounts for the percentage until the channel produces its last read.

From Figures 8, 9, we observe that ReadFish is effective at enriching chromosomes 20 and 21 in this ideal setting of perfect reads and low-latency alignment. Chromosomes 20 and 21 (named "target") are enriched whereas reads coming from other chromosomes are rejected, mostly after 500 bps. Without selective sequencing, the ratio of target to non-target basepairs would be 0.037 ($= \frac{l_{\text{chr20,21}}}{l_{\text{genome}} - l_{\text{chr20,21}}}$ with $l$ being the length), whereas here, we observe roughly 0.58 estimated from the plots, which is a 16-fold improvement. This is very optimistic because we assume perfect basecalling and mapping. The fraction covered is computed by dividing each chromosome into blocks of size `blocksize` (except for the last which is shorter). Whenever a read intersects with a block, we increment that block's count by the number of basepairs intersecting with the read. After adding all reads up to time t, we can compute the fraction of blocks with average coverage at least 1, 2, 3, 4 which is shown in the figure comparing the target to the non-target chromosomes. We use a `blocksize` of 1000 which greatly speeds up computations with respect to `blocksize` 1.

Our script also permits to generate reads with NanoSim and feed them to the simulator. We do not do it here since NanoSim read generation is rather slow. We provide a jupyter script that shows how to generate reads with NanoSim. Note that we fixed bugs in

NanoSim to make runs with the same random seed reproducible and made other small modifications, available as a submodule in our repository, see Section B.1.

### E. Extra Files

The supplemental material contains additional figures from the experiments described here.

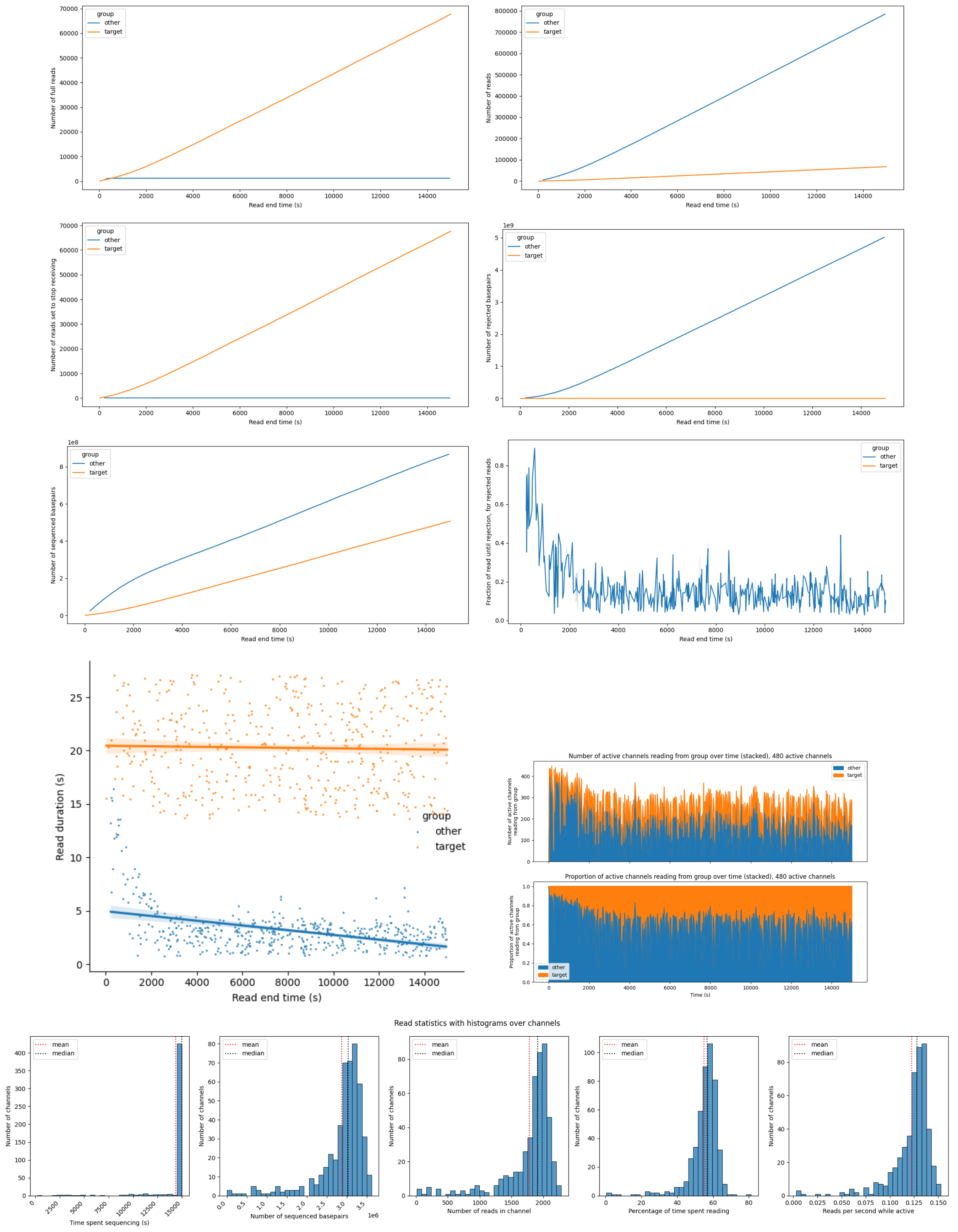


Fig. 8: Combining the simulator with ReadFish to enrich for chromosomes chr20, 21 (target vs other). Number of reads, full reads (reads that were not rejected), reads set to *stop_receiving*, number of rejected and sequenced basepairs, fraction of read until it got rejected, read duration of reads, number and relative proportion of channels reading from target vs other, read stats by channel. The simulation is running for 15000s which is significantly less than the flowcell lifetime, so almost all channels sequence until the end. The read end time of the target group does not start at zero because the reads from the target group do not get rejected, whereas the non-target reads get rejected immediately.

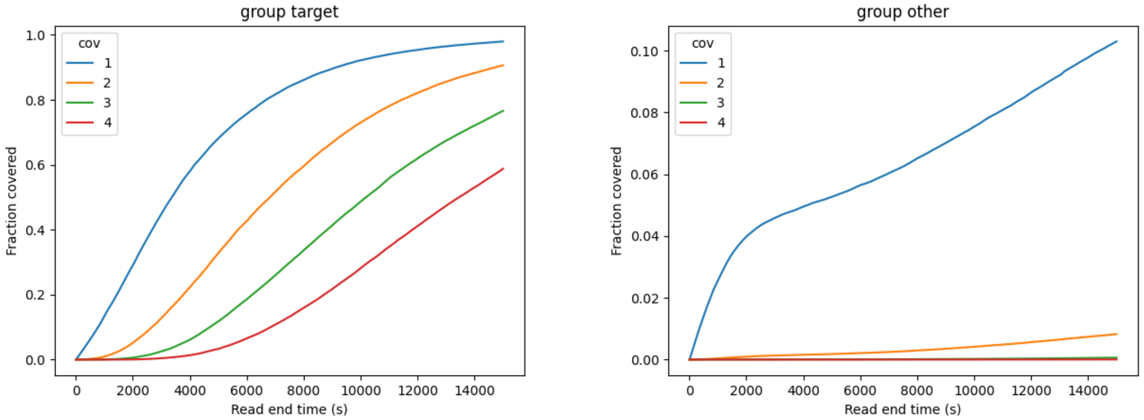


Fig. 9: Combining the simulator with ReadFish to enrich for chromosomes chr20, 21 (target vs other). Fraction of basepairs covered at least $x$ times for $x = 1, 2, 3, 4$ as the sequencing run progresses. Coverage for target is much better than for the rest.
